# Engineered synthetic one-carbon fixation exceeds yield of the Calvin Cycle

**DOI:** 10.1101/2022.10.19.512895

**Authors:** Beau Dronsella, Enrico Orsi, Sara Benito-Vaquerizo, Timo Glatter, Arren Bar-Even, Tobias. J. Erb, Nico J. Claassens

## Abstract

One-carbon (C1) feedstocks derived from CO_2_ and renewable electricity, such as formate, are promising substrates for sustainable production of chemicals, food and fuels. Energetically more efficient, engineered C1-fixation pathways were proposed to increase biomass yields above their natural counterparts, but have so far not been shown to achieve this. Here, we replace the native ‘energy-inefficient’ Calvin-Benson-Bassham (CBB) cycle in *Cupriavidus necator* by genomic integration of the synthetic reductive glycine pathway for growth on formate. Our final engineered strain reaches a higher biomass yield than the CBB-cycle-utilizing wild type, showing for the first time that efficiencies found in natural metabolism can be exceeded via a synthetic pathway. This yield increase demonstrates the potential of synthetic metabolism and is an important step towards realizing truly sustainable, economically feasible bio-based production.

## Main text

Climate change and rising demands for food, fuels and chemicals urgently call to explore alternatives for fossil-based production of carbon-containing molecules. An emerging, promising alternative to plant biomass-based approaches is electromicrobial production^1^. This route consists of efficient electrocatalytic generation of mediator molecules, such as formate, and subsequent microbial conversion of these mediators into biomass or desired products^2^. Recently the biotechnologically relevant microorganisms *Escherichia coli* and *Pichia pastoris* were successfully engineered for formatotrophic growth via transplantation of the CBB cycle ^3^,^4^. Current studies aimed at optimizing the CBB cycle focused on bypassing wasteful photorespiration resulting from the oxygenation side-reaction of the carboxylase Rubisco^5^ or to increase its specificity for CO_2_, while maintaining high turnover^6^. While the latter approach saw limited success so far, a recent reconstruction of Rubisco evolution gave molecular insights that may enable the engineering of improved Rubisco variants in the future^7^. However, apart from limitations related to Rubisco, the inherent ATP inefficiency of the CBB cycle is directly limiting the maximum aerobic yield on formate^8^.

In recent years, alternative more efficient pathway designs were proposed to outcompete the CBB cycle^8–11^, some of which were established *in vitro* and partially implemented *in vivo*^12–14^. However, these efforts have not demonstrated yet that microbial yields on one-carbon-substrates can be surpassed by these synthetic pathways in practice. The most ATP-efficient aerobic pathway for formate and CO_2_ bioconversion is the reductive glycine pathway (rGlyP) (Fig. 1a, S1). This linear pathway requires only 2 ATP to generate one pyruvate molecule, unlike the CBB cycle, which requires 7 ATP.

**Fig. 1:**
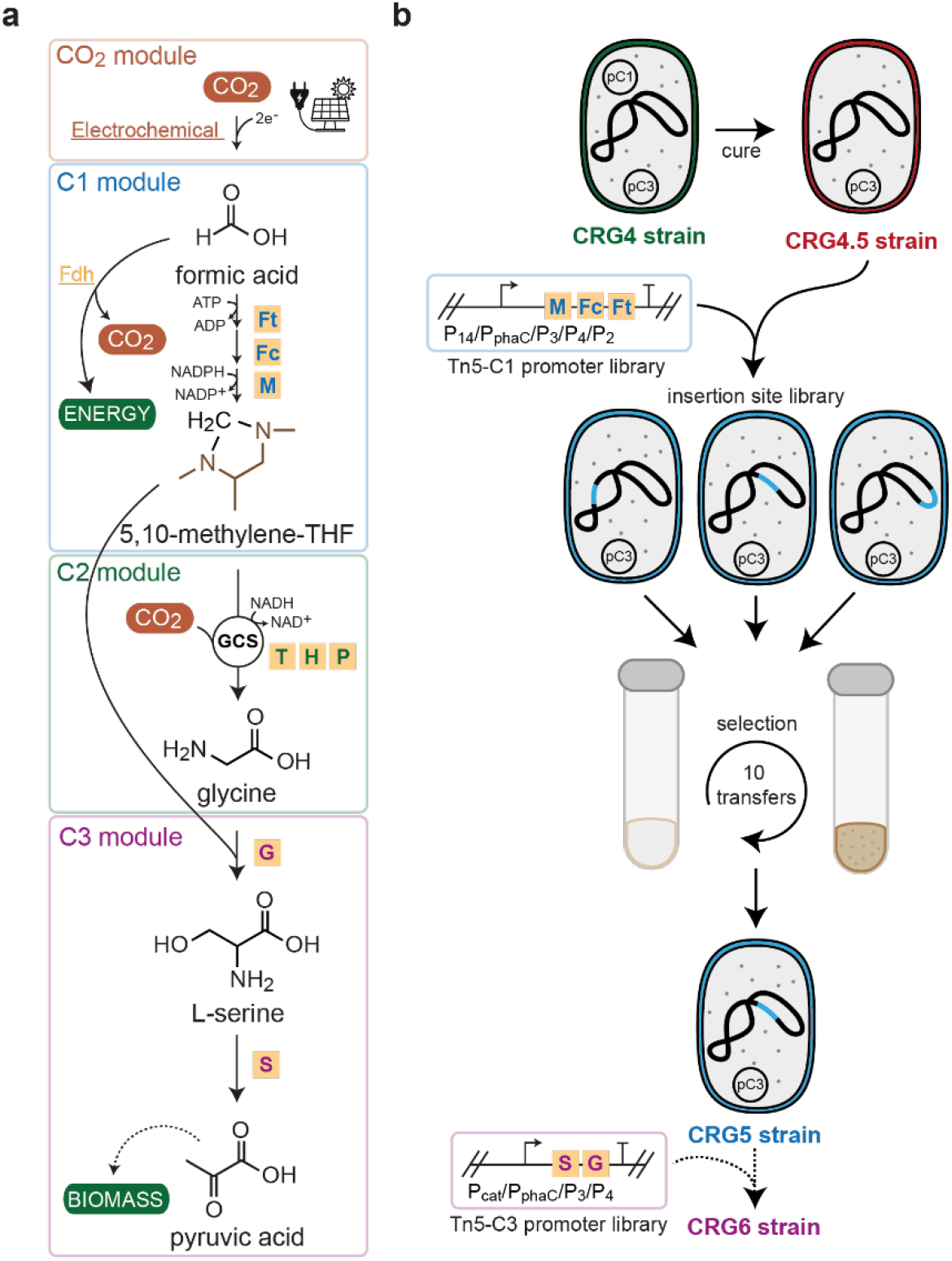
The reductive glycine pathway and the strategy of genomic pathway integration. **a**, Architecture of the reductive glycine pathway (rGlyP) with the proposed serine route as option to metabolize glycine. In the CO_2_ module, formate is generated from electrochemical reduction of CO_2_. The C1 module then activates and reduces formate to 5,10-methylene-THF via the 3 heterologous enzymes from *M. extorquens* formate-THF ligase (FtfL, Ft), 5,10-methenyl-THF-cyclohydrolase (FchA, Fc) and 5,10-methylene-THF dehydrogenase (MtdA, M). Next methylene-THF is converted to glycine by the glycine cleavage system (gcvTHP, THP) operating in the reductive direction (C2 module). Last, glycine is condensed with another methylene-THF via serine hydroxymethyltransferase (GlyA, G) to yield serine, which is then dehydrated by serine deaminase (SdaA, S) to pyruvate. Energy is generated from formate via formate dehydrogenase. Formate and pyruvate are referred to as such in the text for consistency sake, but are here depicted in their protonated forms, in which they are actually metabolized. **b**, *C. necator* strain engineering workflow employed in this study. The operons previously expressed from plasmids were consecutively genomically integrated using a combined approach of selection for pathway activity and Tn5-transposon-mediated integration.

While the rGlyP was conceptualized as a synthetic, aerobic pathway for formate and CO_2_ fixation, an anaerobic naturally existing variant of this pathway was recently discovered in the anaerobic bacterium *Desulfovibrio desulfuricans*^15^. Before its discovery in nature, however, the rGlyP was already successfully engineered via a modular and laboratory evolution approach for full formatotrophic growth in different model organisms, including *Cupriavidus necator^16–18^. C. necator* is an aerobic formatotroph that natively utilizes the CBB cycle for chemolithoautotrophic growth. Moreover, *C. necator* is a key organism for electromicrobial production that was utilized in several proof-of-principle studies for electricity-based production from CO_2_^19,20^. One of these studies even indicated that electromicrobial production could outcompete the solar-to-product efficiency of biological photosynthesis^20^. *C. necator* is also a prime candidate for electromicrobial production of food and feed, which can potentially result in many-fold higher aerial protein yields than current agricultural protein production^21,22^. Electromicrobial production could be even further increased if higher formatotrophic yields were obtained, which could potentially be realized via the rGlyP. However, despite the theoretical ATP advantage of the rGlyP, our previously engineered and evolved *C. necator* strain harboring the full rGlyP (CRG4) did not exceed the formatotrophic biomass yield of the original CBB cycle-harboring strain^18^.

Here, we engineer a *C. necator* strain expressing the full rGlyP from the genome that exceeds the biomass yield of the CBB-cycle-harboring wild-type strain on formate. To achieve this we employ a recently updated genome-scale metabolic model of *C. necator^23^* to predict the rGlyP to outcompete the formatotrophic yield of the CBB cycle by ~17 % (Fig. S2). Since most of the rGlyP is expressed from plasmids in CRG4 (Fig. 1b), we reason that plasmid replication elements and/or unbalanced (over)expression of pathway genes may cause burden to the strain, lowering the final biomass yield. Therefore, we employ a Tn5-transposon-based approach^24^ for the stepwise genomic integration of the full rGlyP coupled with selection (and/or evolution) for growth on formate (Fig. 1b). Last, we use proteomics to elucidate the cellular adaptations that enable our genomic rGlyP strain to surpass its predecessors and the CBB-cycle-harboring strain.

In a first step, we cured the CRG4 strain^18^ of the plasmid expressing the first module of the rGlyP (pC1 in Fig. 1b). Since this strain already contained a synthetic constitutive promoter expressing the C2 module, it was an ideal platform to select for C1 module activity on formate (Fig. S3c). Exploiting this, we then integrated the C1 module into the genome using the Tn5 transposon machinery and subsequent selection for activity of the rGlyP via growth on formate (Supplementary Text S1, Fig. S4a, S5a). This fast-growing strain with a genomically expressed module 1 (CRG5) was then characterized and compared to the CRG4 and CBB strains for growth and biomass yield formation on formate. Genomic integration of the C1 module resulted in faster growth compared to CRG4, albeit still slower than the wild-type strain harboring the CBB Cycle (Fig. 2a.). CRG5 also reached a higher maximal OD_600_ compared to both CRG4, as well as the wild-type strain (Fig. S3a). This demonstrated for the first time that the formatotrophic biomass yield of the rGlyP might exceed the yield of the CBB Cycle in *C. necator*. Sequencing of CRG5 revealed two insertions of the C1 module, one in the gene encoding 2-methyl-cis-aconitate hydratase (*acnM*) and another in the *phaP1* promoter locus (Table S1, Fig. S6a). We hypothesized that the transposon insertion strategy achieved two things simultaneously. First, it downregulated the abundant phasin protein PhaP1, which is not required in the non-PHB-producing Δ*phaC* strain background. Second, expression of the integrated C1 module benefitted from a read-through effect from the adjacent *phaP1* promoter.

**Fig. 2:**
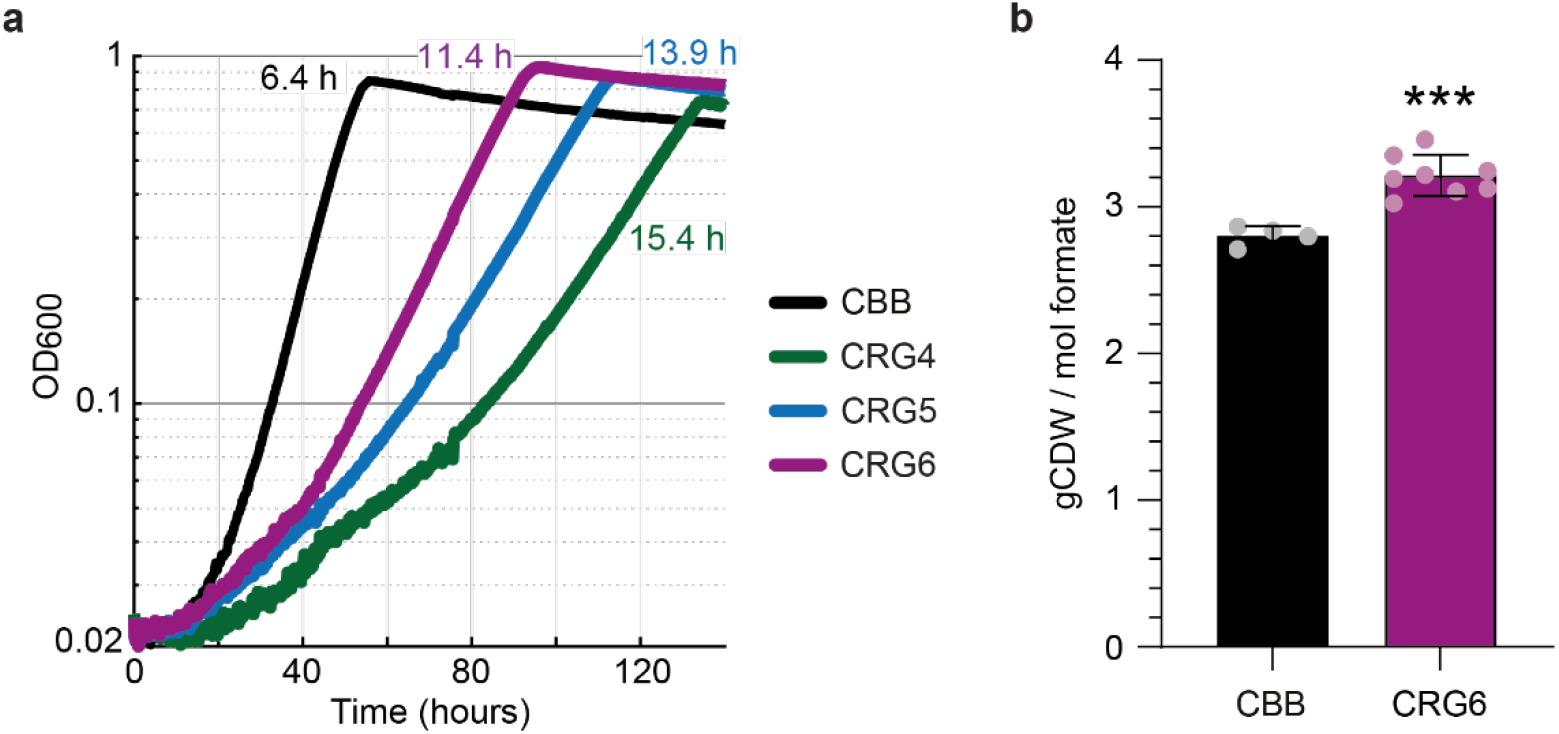
Characterization of *C. necator* strains growing via the rGlyP compared to the CBB cycle. **a**, Growth of *C. necator H16* Δ*phaC* strains harboring the CBB cycle compared to CRG4, CRG5 and CRG6 in M9 minimal media supplemented with 80 mM formate and 100 mM bicarbonate with 10 % CO_2_ in the headspace. The doubling time in hours (h) of the strains is presented in the designated strain color. Curves depict the mean of at least 2 technical replicates and are representative of 3 experiments conducted in the same conditions to ensure reproducibility. **b,** Biomass yields in gram cell dry weight per mol formate consumed of CBB and CRG6 strains grown in formate minimal media in shake flasks. Data were obtained from 4 and 8 biological replicates of 2 and 3 experiments respectively (n ≥ 4). All data points and their mean are shown. Error bars indicate standard deviation. Three asterisks (***) represent a p-value < 0.001.

To improve the strain further, we next integrated also the C3 module into the genome of CRG5 (cured of pC3) using the same approach as above (Supplementary Text S2, Fig. S4b, S5b, S6b). This CRG6 strain now harbored a fully genomically integrated rGlyP and grew even faster than CRG4 or CRG5 at a doubling time of ~11 hours, which was (still) slower than the wild-type CBB strain (~6 hours doubling time, Fig. 2a). However, CRG6 consistently grew to a higher maximum OD_600_ than both CRG4 and, most importantly, the CBB cycle strains (Fig. 2a, S3a). To confirm that more biomass was formed by CRG6, we determined the biomass yield of CRG6 and wild-type CBB cycle strain on formate minimal media. This revealed that both CRG6 (and CRG5) formed about 14 % more biomass per mol of formate compared to the CBB cycle strain (3.2 and 2.8 gCDW/mol formate, respectively) coming close to the theoretically predicted maximum yield increase for the rGlyP of 17 % (Fig. 2b, S2, S3b).

To investigate the cellular adaptations causing the yield increase of CRG6, we performed label-free quantitative proteomics for relative comparison and total abundance estimation of the proteomes of CRG6, CRG4 and CBB strains (Supplementary Text S3, Fig. 3a,b, S7a). This demonstrated that genomic integration of the C1 module resulted in similar expression levels in CRG6 compared to plasmid-based expression in CRG4, indicating that indeed a read-through effect from the *phaP1* promoter had occurred (Fig. 3a). The C1-operon insertion further led to a 150-fold downregulation of PhaP1 resulting in a decrease of its quantified proteome fraction from ~1 % to ~0.01 % in CRG4 versus CRG6 (see Source data). The protein levels of the genomic C3 module in CRG6 were reduced more than 10-fold compared to CRG4, suggesting that this module was highly overexpressed in CRG4 (Fig. 3b). In addition, plasmid backbone protein expression (~0.4 % of quantified proteome in CRG4), possibly requiring additional energy investment for plasmid replication, was completely absent in CRG6 (Fig. 3b). This loss of plasmid burden and maintenance together with reduction of proteome allocation to the non-functional PHB biosynthesis machinery might have caused reduced energy consumption and may explain the increased yields observed for CRG6.

**Fig. 3:**
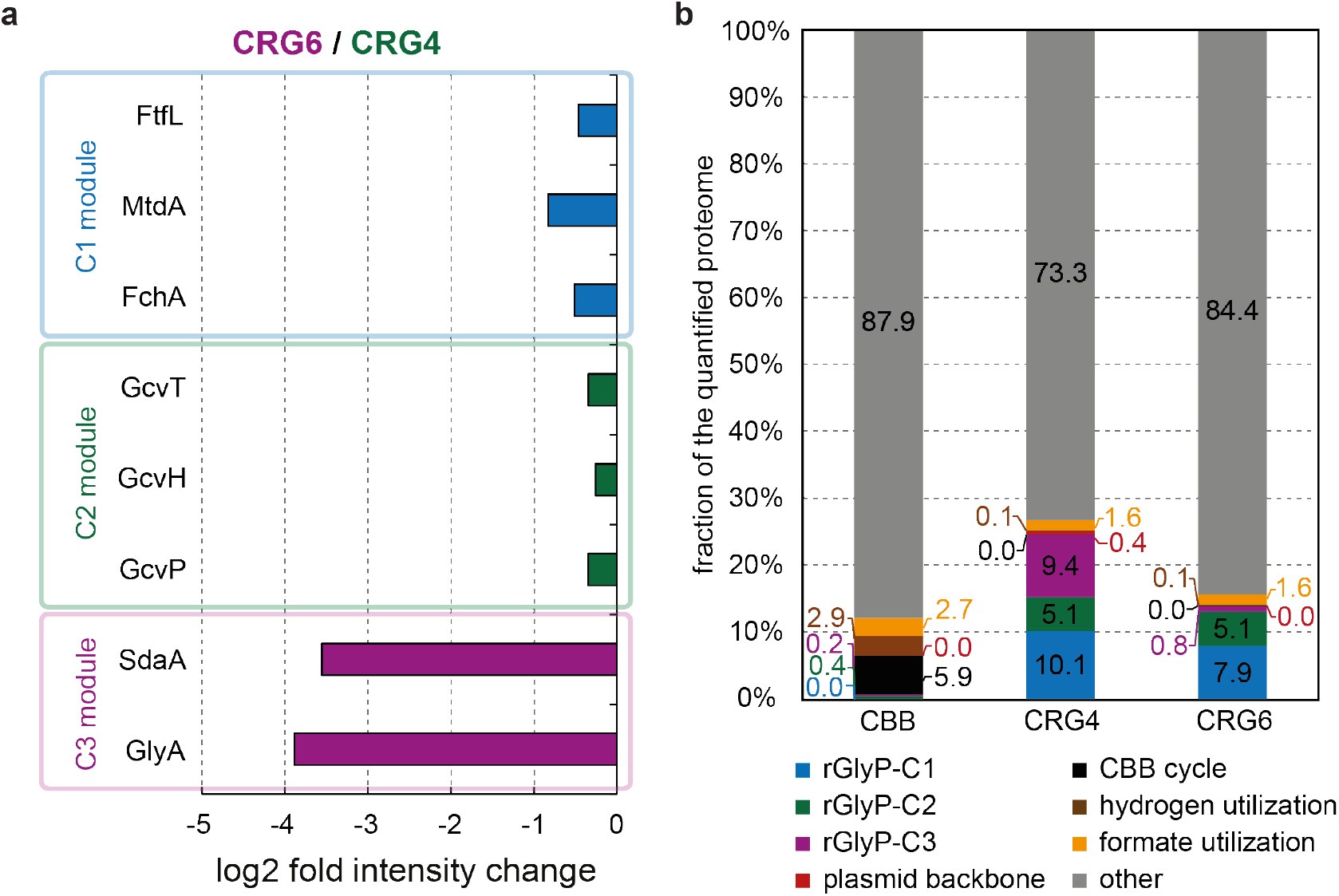
Proteomic comparison of rGlyP and CBB strains. **a**, Relative protein intensity changes of the rGlyP modules in strain CRG6 relative to CRG4 both grown on formate minimal media. **b**, Fractions of the quantified proteome associated with various metabolic tasks in the strains CBB, CRG4 and CRG6 all grown on formate minimal media. Clustering criteria for grouping of proteins by metabolic tasks are provided in the methods section.

Last we turned our attention to another key enzyme for operation of both the rGlyP and CBB cycle, i.e., the energy-generating formate dehydrogenase (FDH). *C. necator* natively harbors a molybdenum-dependent NAD-reducing soluble FDH (sFDH) with catalytic activities more than two orders of magnitude higher than non-metal-dependent FDHs^25^,^26^. This is in stark contrast to the published formatotrophic *E. coli* rGlyP strains that were dependent on a strongly expressed heterologous non-metal-dependent FDH for fast growth^16^. Note that while the CBB cycle requires all formate to be oxidized, the rGlyP depends on a balancing of flux between formate assimilation and formate oxidation. To investigate if biomass yields could be increased even further, we performed tungstate titration experiments to inhibit sFDH activity^26^. However, no further increases in biomass yield could be obtained over a range of almost 6 orders of magnitude of supplemented tungstate. This indicated that formate over-oxidization via sFDH in particular was not limiting the biomass yield in strain CRG6 (Supplementary Text S4, Fig. S8). Instead, our proteomics data revealed that the total FDH fraction was decreased by almost half between CBB cycle strain (~2.7 % of quantified proteome) and CRG4/6 (~1.6 %) (Fig. 3b, S7b,c), possibly reflecting a different solution to balance oxidative and assimilatory formate flux.

Overall, this work shows for the first time that an engineered C1-assimilation pathway can outcompete the biomass yield of a natural pathway in the same organism. This also provides evidence that other designed pathways with theoretically higher efficiencies can improve *in vivo* yields. In the future, these promising synthetic one-carbon fixation pathways, which were only demonstrated *in vitro* so far, e.g., the CETCH cycle for CO_2_ fixation^10,12^, may outperform the CBB cycle *in vivo*, once successfully established. The (randomized) transposon integration technique and growth-coupled selection approach in this work could serve as a blueprint to realize efficient pathway implementation in more difficult-to-engineer bacteria, as well as eukaryotes such as yeast and microalgae. This could enable the engineering of superior one-carbon assimilation performance in these hosts. Realizing higher yields and faster growth on one-carbon substrates will aid to make the electromicrobial production route an attractive, economically feasible reality for sustainable production of carbon-based chemicals, fuels, feed and food.

## Acknowledgements

We dedicate this work to the memory of Arren Bar-Even (1980-2020) who was involved in the design and the initial phase of this study. Above all he has been an exceptional and dear mentor and colleague to the authors. We like to thank Oliver Lenz for providing plasmid pLO3 and advice on working with *Cupriavidus necator*. In addition, we acknowledge Pablo Nikel for providing the pBAMD plasmids and William Newell and Joerg Kahnt for experimental assistance. We thank Michael Jahn for input on the quantitative proteomic data analysis and Georgia Angelidou for help with bioinformatic analysis. Last, we thank Helena Schulz-Mirbach, Ari Satanowski and Sebastian Wenk for critical reading of the manuscript. B.D. and E.O. are financially supported by the German Ministry of Education and Research (BMBF) through the grant Transformate (033RC023G) and the Max Plank Society. N.J.C. acknowledges support from a Veni grant (VI.Veni.192.156) from the Dutch Science Organization (NWO).

## Author contributions

A.B-E., B.D. E.O, and N.J.C. designed and conceived the study. A.B-E., N.J.C. and T.J.E. supervised the project. B.D. performed the strain engineering, experimental characterization and NGS analysis. T.G and B.D performed the proteomic analysis. S.B.V performed modelling. B.D., E.O., T.J.E. and N.J.C. wrote the manuscript. All others provided feedback and approved the manuscript.

## Supplementary material

### Materials and Methods

#### Bacterial strains and conjugation

*C. necator H16* deleted in polyhydroxybutyrate biosynthesis (Δ*phaC1*) served as the base strain (‘wild type’) in this work. All CRG strains are also deleted for carbon fixation via the Calvin cycle by deletion of both Rubisco subunits on chromosome 2 and megaplasmid (Δ*cbbSL2, ΔcbbSLp*). Routine cloning was performed in *E. coli* DH5α, while *E. coli* ST18 cells were used for conjugation of mobilizable plasmids to *C. necator* via biparental spot mating.

#### Cultivation conditions

*C. necator* and *E. coli* were grown on Lysogeny Broth (LB) (10 g/L NaCl, 5 g/L yeast extract and 10 g/L tryptone) for routine cultivation and genetic modifications. When appropriate the antibiotics kanamycin (100 μg/mL for *C. necator* and 50 μg/mL for *E. coli*), chloramphenicol (30 μg/mL), tetracycline (10 μg/ mL), ampicillin (100 μg/mL for *E. coli*) or gentamycin (20 μg/mL for *C. necator*) were added. Growth experiments were conducted in M9 minimal medium (47.8 mM Na_2_HPO_4_, 22 mM KH_2_PO_4_, 8.6 mM NaCl, 18.7 mM NH_4_Cl, 2 mM MgSO_4_ and 100 μM CaCl_2_), supplemented with trace elements (134 μM EDTA, 31 μM FeCl_3_·6H_2_O, 6.2 μM ZnCl_2_, 0.76 μM CuCl_2_·2H_2_O, 0.42 μM CoCl_2_·2H_2_O, 1.62 μM H_3_BO_3_, 0.081 μM MnCl_2_·4H_2_O). Routine cultivation was performed in 4 mL medium in 12 mL glass tubes in an orbital shaker incubator at 240 rpm at 30°C and 37°C for *C. necator* and *E. coli* respectively. M9 minimal media was supplemented with 80 mM sodium formate, 100 mM sodium bicarbonate, pH adjusted to 7.2, under a headspace of 10 % CO_2_ (v/v) for formatotrophic growth. No antibiotics were added during growth characterization experiments in the plate reader. Growth measurements were obtained from 96 well-plate experiments (Nunc transparent flat-bottom, Thermo Scientific). Strains were typically pre-cultured in M9 minimal medium supplemented with 20 mM pyruvate. Cells were harvested, washed twice and inoculated at an OD_600_ of 0.01. 150 μl of cell medium mix was topped with 50 μl transparent mineral oil (Sigma-Aldrich) to avoid evaporation, while maintaining diffusion of O_2_ and CO_2_. 96-well plates were incubated at 30°C with continuous shaking (alternating between 30 sec orbital and 30 sec linear) in a Tecan infinite M200Pro plate reader. OD_600_ values were measured every 8 min. Growth data were blanked and converted from plate reader OD_600_ to cuvette OD_600_ by multiplication with a factor of 4.35 via a Matlab script. All growth experiments were repeated at least three times, and the growth curves shown are representative curves of these experiments.

**Table 1:** A complete overview of strains used in this study. Plasmid-based (p) or genomic (g) expression of the genes 5,10-methylene-THF dehydrogenase (*mtdA*, UniProt: P55818), 5,10-methenyl-THF cyclohydrolase (*fchA*, UniProt: Q49135) and formate-THF ligase (*ftfL*, UniProt: Q83WS0) from *Methylorubrum extorquens AM1, gcvTHP, sdaA* and *glyA* from *C. necator* under the control of different promoters.

| Strain | Genotype | Source |
| --- | --- | --- |
| *E. coli* DH5α | F- λ- Φ80*lacZ*ΔM15 Δ(*lacZYA-argF*)U169 *deoR*<br>*recA1 endA1*<br>*hsdR17*(rK- mK+) *phoA supE44 thi-1 gyrA96*<br>*relA1* | Lab stock |
| *E. coli* ST18 | *pro thi hsdR+* Tpr Smr; chromosome::RP4-2<br>Tc::Mu-Kan::Tn7/λpir<br>Δ*hemA* | Lab stock |
| *C. necator* H16 | Wild type | Lab collection |
| Δ*phaC1* | Δ*phaC1* | Lab collection |
| ΔCBB | Δ*phaC1*, Δ*cbbSL2p* | Claassens 2020 |
| CRG4 | ΔCBB +pSEVA221-P<sub>14</sub>-*mtdA-fch-fftL* +gP<sub>3mut2</sub>-*gcvTHP* +pSEVA331-P<sub>phaC1</sub>-*sdaA-glyA* | Claassens 2020 |
| CRG4.5 | ΔCBB +gP<sub>3mut2</sub>-*gcvTHP* +pSEVA331-P<sub>phaC1</sub>-*sdaA-glyA* | This study |
| CRG5 | ΔCBB +gP<sub>14</sub>-*mtdA-fch-fftL*-Tn5 +gP<sub>3mut2</sub>-*gcvTHP* +pSEVA331-P<sub>phaC1</sub>-*sdaA-glyA* | This study |
| CRG5-P<sub>Phac</sub>-C1 | ΔCBB +gP<sub>Phac</sub>-*mtdA-fch-fftL*-Tn5 +gP<sub>3mut2</sub>-*gcvTHP* +pSEVA331-P<sub>phaC1</sub>-*sdaA-glyA* | This study |
| CRG5-P<sub>3</sub>-C1 | ΔCBB +gP<sub>3</sub>-*mtdA-fch-fftL*-Tn5 +gP<sub>3mut2</sub>-*gcvTHP* +pSEVA331-P<sub>phaC1</sub>-*sdaA-glyA* | This study |
| CRG5-P<sub>4</sub>-C1 | ΔCBB +gP<sub>4</sub>-*mtdA-fch-fftL*-Tn5 +gP<sub>3mut2</sub>-*gcvTHP* +pSEVA331-P<sub>phaC1</sub>-*sdaA-glyA* | This study |
| CRG5-P<sub>2</sub>-C1 | ΔCBB +gP<sub>2</sub>-*mtdA-fch-fftL*-Tn5 +gP<sub>3mut2</sub>-*gcvTHP* +pSEVA331-P<sub>phaC1</sub>-*sdaA-glyA* | This study |
| CRG5-P<sub>14</sub>-C1-2 | ΔCBB +gP<sub>14</sub>-*mtdA-fch-fftL*-Tn5 +gP<sub>3mut2</sub>-*gcvTHP* +pSEVA331-P<sub>phaC1</sub>-*sdaA-glyA* | This study |
| CRG5-P<sub>Phac</sub>-C1-2 | ΔCBB +gP<sub>Phac</sub>-*mtdA-fch-fftL*-Tn5 +gP<sub>3mut2</sub>-*gcvTHP* +pSEVA331-P<sub>phaC1</sub>-*sdaA-glyA* | This study |
| CRG5-P<sub>3</sub>-C1-2 | ΔCBB +gP<sub>3</sub>-*mtdA-fch-fftL*-Tn5 +gP<sub>3mut2</sub>-*gcvTHP* +pSEVA331-P<sub>phaC1</sub>-*sdaA-glyA* | This study |
| CRG5-P<sub>4</sub>-C1-2 | ΔCBB +gP<sub>4</sub>-*mtdA-fch-fftL*-Tn5 +gP<sub>3mut2</sub>-*gcvTHP* +pSEVA331-P<sub>phaC1</sub>-*sdaA-glyA* | This study |
| CRG5-P<sub>2</sub>-C1-2 | ΔCBB +gP<sub>2</sub>-*mtdA-fch-fftL*-Tn5 +gP<sub>3mut2</sub>-*gcvTHP* +pSEVA331-P<sub>phaC1</sub>-*sdaA-glyA* | This study |
| CRG5.5 | ΔCBB +gP<sub>14</sub>-*mtdA-fch-fftL*-Tn5 +gP<sub>3mut2</sub>-*gcvTHP* | This study |
| CRG6 | ΔCBB +gP<sub>14</sub>-*mtdA-fch-fftL*-Tn5 +gP<sub>3mut2</sub>-*gcvTHP* +gP<sub>3</sub>-*sdaA-glyA*-Tn5 | This study |
| CRG6-P<sub>cat</sub>-C3 | ΔCBB +gP<sub>14</sub>-*mtdA-fch-fftL*-Tn5 +gP<sub>3mut2</sub>-*gcvTHP* +gP<sub>cat</sub>-*sdaA-glyA*-Tn5 | This study |
| CRG6-P<sub>Phac</sub>-C3 | ΔCBB +gP<sub>14</sub>-*mtdA-fch-fftL*-Tn5 +gP<sub>3mut2</sub>-*gcvTHP* +gP<sub>Phac</sub>-*sdaA-glyA*-Tn5 | This study |
| CRG6-P<sub>4</sub>-C3 | ΔCBB +gP<sub>14</sub>-*mtdA-fch-fftL*-Tn5 +gP<sub>3mut2</sub>-*gcvTHP* +gP<sub>4</sub>-*sdaA-glyA*-Tn5 | This study |
| CRG6-P<sub>3</sub>-C3-2 | ΔCBB +gP<sub>14</sub>-*mtdA-fch-fftL*-Tn5 +gP<sub>3mut2</sub>-*gcvTHP* +gP<sub>3</sub>-*sdaA-glyA*-Tn5 | This study |
| CRG6-P<sub>cat</sub>-C3-2 | ΔCBB +gP<sub>14</sub>-*mtdA-fch-fftL*-Tn5 +gP<sub>3mut2</sub>-*gcvTHP* +gP<sub>cat</sub>-*sdaA-glyA*-Tn5 | This study |
| CRG6-P<sub>Phac</sub>-C3-2 | ΔCBB +gP<sub>14</sub>-*mtdA-fch-fftL*-Tn5 +gP<sub>3mut2</sub>-*gcvTHP* +gP<sub>Phac</sub>-*sdaA-glyA*-Tn5 | This study |
| CRG6-P<sub>4</sub>-C3-2 | $\Delta CBB$ +gP<sub>14</sub>-*mtdA-fch-fftL*-Tn5 +gP<sub>3mut2</sub>-*gcvTHP* +gP<sub>4</sub>-*sdaA-glyA*-Tn5 | This study |
| CRG6 $\Delta dadA6$ | $\Delta CBB$, $\Delta dadA6$ +gP<sub>14</sub>-*mtdA-fch-fftL*-Tn5 +gP<sub>3mut2</sub>-*gcvTHP* +gP<sub>3</sub>-*sdaA-glyA*-Tn5 | This study |

**Table 2:** A complete overview of plasmids used in this study.

| Abbreviation | Name | Source |
| --- | --- | --- |
| pC1 | pSEVA221-P<sub>14</sub>-*mtdA-fch-fftL* | Claassens 2020 |
| pC3 | pSEVA331-P<sub>PhaC1</sub>-*sdaA-glyA* | Claassens 2020 |
| pBAMD-P<sub>14</sub>-C1 | pBAMD1-4-P<sub>14</sub>-*mtdA-fch-fftL* | This study |
| pBAMD-P<sub>PhaC</sub>-C1 | pBAMD1-4-P<sub>PhaC</sub>-*mtdA-fch-fftL* | This study |
| pBAMD-P<sub>3</sub>-C1 | pBAMD1-4-P<sub>3</sub>-*mtdA-fch-fftL* | This study |
| pBAMD-P<sub>4</sub>-C1 | pBAMD1-4-P<sub>4</sub>-*mtdA-fch-fftL* | This study |
| pBAMD-P<sub>2</sub>-C1 | pBAMD1-4-P<sub>2</sub>-*mtdA-fch-fftL* | This study |
| pBAMD-P<sub>cat</sub>-C3 | pBAMD1-4-P<sub>3</sub>-*sdaA-glyA* | This study |
| pBAMD-P<sub>PhaC</sub>-C3 | pBAMD1-4-P<sub>3</sub>-*sdaA-glyA* | This study |
| pBAMD-P<sub>3</sub>-C3 | pBAMD1-4-P<sub>3</sub>-*sdaA-glyA* | This study |
| pBAMD-P<sub>4</sub>-C3 | pBAMD1-4-P<sub>3</sub>-*sdaA-glyA* | This study |
| pLO3-dadA6 | pLO3-dadA6 | Claassens 2020 |

**Table 3:**
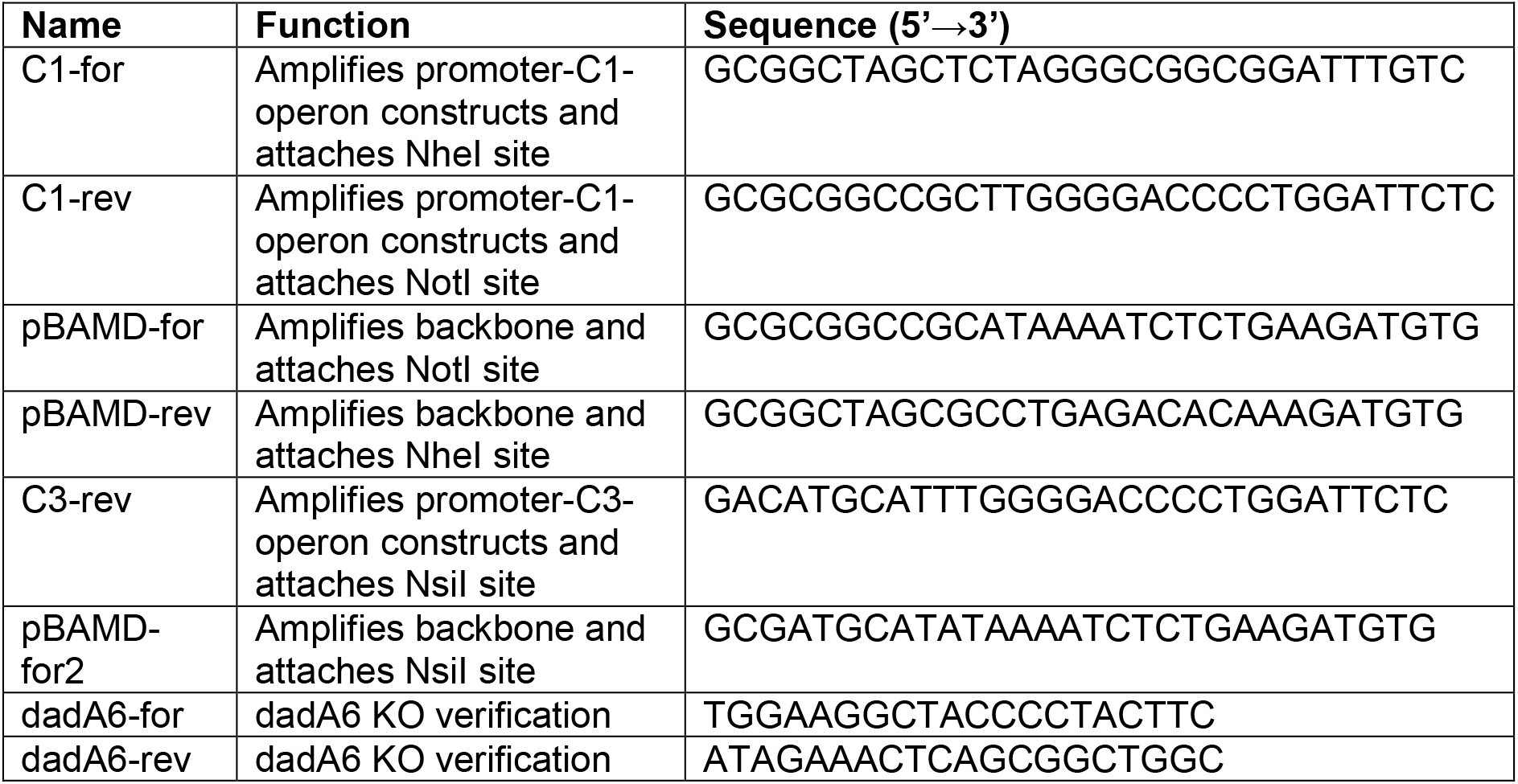
DNA oligo primers used in this study.

#### Tn5 vector construction

The vector pBAMD1-4 was kindly provided by Pablo I. Nikel. The vector backbone was amplified with primers pBAMD-for and pBAMD-rev without the antibiotic resistance gene in the cargo module (module between the transposon recognition sites ME1 and ME2) and NotI and NheI restriction sites were attached. The operons *P_14_/P_PhaC_/P_3_/P_4_/P_2_-mtdA-fch-ftfL* and Pcat/P_PhaC_/P_3_/P_4_-*sdaA*-*glyA* were amplified from existing pSEVA221 and pSEVA331 based expression plasmids respectively with primers C1-for and C1-rev attaching NotI and NheI restrictions sites. Promoter-FCM/SG operons were cloned into pBAMD vectors using restriction ligation.

All genes of the C1 module were previously placed behind a synthetic RBS designed with RBS Calculator with a translation initiation rate of 30.000 arbitrary units^27^.

#### DadA6 knockout

The plasmid pLO3-dadA6 was utilized to delete the *dadA6* gene in the CRG6 strain via allelic replacement based on sucrose counter selection with SacB as described previously^18^,^28^. The deletion was confirmed by PCR using the primers dadA6-for and dadA6-rev and whole genome sequencing.

#### Tn5 mediated knock-in coupled with liquid selective conditions

*E. coli* ST18 strains harboring the P_14_/P_PhaC_/P_3_/P_4_/P_2_-C1 and Pcat/P_PhaC_/P_3_/P_4_-C3 constructs in pBAMD-Tn5 vectors served as donor strains. *C. necator* strains CRG4.5 and CRG5.5 served as recipient strains for the conjugation (Fig. S3c). 100μl of LB grown dense overnight cultures of *C. necator* and *E. coli* ST18 (supplemented with 50 μg/mL 5-amniolevulinic acid (ALA) were mixed in a 1.5 mL plastic tube. 100 μl of this cell mixture was plated on LB+ALA agar plates and dried for 30 min before incubating overnight at 30°C. The next day the grown cell lawn was resuspended in LB and a 100 μl inoculum of an OD_600_=1 mixture of *C. necator* recipient and *E. coli* ST18 donor cells was used to inoculate 4 mL liquid M9 minimal media supplemented with 80 mM formate, 100 mM bicarbonate in 10 % CO_2_ (v/v). Upon observing cellular growth and the population reaching late log to stationary phase, 1 μL of OD_600_=1 culture was used to re-inoculate into 4 mL of selective media. This population was passaged 10 times to select for better growth (Fig. S4). After passage 10 the population was dilution streaked two consecutive times to single colony on LB agar plates with 20 μg/mL gentamicin. The isolated single clones were compared to the populations for the growth behavior in the selective formate media and saved in 25 % glycerol at −80°C (Fig. S5).

#### Biomass yield experiments and determination of formate concentration

For the determination of cell dry weight (CDW) of *C. necator* grown on formate 250 mL Pyrex Erlenmeyer flasks with 50 mL M9 media supplemented with 80 mM formate and 100 mM bicarbonate were inoculated to a starting OD_600_ of 0.002 from cells pre-cultured in the same media in glass tubes. The flasks were incubated in an Infors Minitron at 30°C in an atmosphere of 10 % CO_2_ (v/v) at 240 rpm. 50 mL of formate grown culture at the end of the exponential phase (OD_600_ 0.6-0.9) was harvested by centrifugation at 3220 *g* for 20min and washed 3 times in ddH_2_O to remove residual media components. Washed cells were then pipetted into custom-made pre-dried and pre-weighed aluminum trays. The samples were dried for a period of 24h at 90°C and were then weighed to obtain the additional weight from the dry cells. The leftover formate concentrations from the supernatant of the same cultures were measured via a Dionex ICS 6000 HPAEC Ion Chromatography (IC) system from Thermo Fischer, equipped with an organic acid column (Dionex IonPac AS11-HC-4μm) at 30 °C with a flow of 0.38 mL/min using ddH_2_O as eluent. A standard series of M9 minimal media with the formate concentrations (0, 16, 24, 36, 54, 80 mM) was measured in triplicates and their averages were used to calculate the slope of the standard curve for each experiment. The samples and standards were diluted 1:100 before measuring. The biomass yield in gram CDW per mol formate was then calculated by dividing the CDW concentration (g/L) by the decrease in formate concentration (mol/L).

#### Whole-genome sequencing

*C. necator* genomes and plasmids were extracted from LB grown cells for whole-genome sequencing using the NucleoSpin® Microbial DNA Kit (Macherey-Nagel, Düren, Germany). Samples were sent for library preparation (Nextera, Ilumina) and sequencing at an Ilumina NovoSeq 6000 platform to obtain 150bp paired-end reads (Novogene, Cambridge, UK). Samples were paired, trimmed and assembled to the *C. necator* reference genome using the Geneious 8.1 software or the Breseq pipeline. Mutations (frequency above >60 %) were identified based on comparative analysis with the parental strains.

#### Statistics

Statistical analysis of the cell dry weight data was conducted using Graph Pad Prism. One-way ANOVA with multiple comparisons was used to determine significant differences of the means of the CBB-cycle-harboring wild type, CRG4, CRG5, CRG5.5, CRG6 and CRG6 Δ*dadA6* relative to the CBB (Δ*phaC*) group.

#### Constraint-based metabolic modelling

The genome-scale metabolic model (GEM) of *C. necator*, RehMBEL1391_sbml_L3V1 was retrieved in SBML format level 3 version 1 from the public repository provided by Jahn et al^23^. Model simulations were performed using COBRApy 0.24.0^29^ and Python 3.9.

Flux balance analysis (FBA)^30^ was implemented to simulate formate assimilation for both the CBB cycle and the rGlyP. To simulate formate assimilation through the CBB cycle, the glycine cleavage system reaction (“GLYAMT”) was set as non-reversible to prevent any flux through the rGlyP. To support formate assimilation through rGlyP, two additional reactions were added to the GEM: Formate tetrahydrofolate ligase (“Ftl”) and Methenyltetrahydrofolate cyclohydrolase (“Fch”). Additionally, GLYAMT reaction was set as reversible, and the flux through the Ribulose-bisphosphate carboxylase reaction (“RBPC”) was set to 0 to prevent flux through the CBB cycle.

The ATP hydrolysis part of the biomass synthesis reaction was amended with a H_2_O molecule and H^+^ to balance the reaction.

To fit GAM/NGAM values, we iteratively calculated biomass yields for each GAM/NGAM combination selected. GAM was varied between 0 to 400 mmol gDW^-1^, and NGAM was varied between 0 to 8.5 mmol gDW ^-1^ h^-1^. For each GAM value, the biomass synthesis reaction was updated and was constrained to the growth rate of 4 hours and specified NGAM. The formate uptake rate reaction (“EX_formate_e”) was used as the objective function. The ratio between growth rate (h^-1^) and the maximum predicted formate uptake rate (mmol gDW ^-1^ h^-1^) was used to calculate the biomass yields in gDW mol^-1^ formate.

#### Proteomic analysis

*C. necator* CBB, CRG4 and CRG6 strains were pre-cultured in 4 mL M9, minimal media supplemented with 80 mM formate and 100 mM HCO_3_ in 15 mL glass tubes. 1 mL of cells in late exponential phase were harvested and washed 3 times in M9 media without carbon source. From here 50 mL of M9, minimal media supplemented with 80 mM formate and 100 mM HCO_3_ was inoculated to a starting OD_600_ of 0.01 in 250 mL non-baffled shake flasks. Cultures were incubated in an Infors Minitron at 30°C in an atmosphere of 10 % CO_2_ (v/v) at 200 rpm. This flask adapted culture was then used to inoculate a second flask culture (the third consecutive formate cultivation) in the same way. Cells from 3 biological replicates were harvested in mid log phase (OD_600_=0.3-0.5) and washed twice with phosphate buffer (12 mM phosphate buffer, 2.7 mM KCl, 137 mM NaCl, pH=7.4). Cell pellets corresponding to and OD_600_ of 3 were flash frozen in liquid nitrogen and stored at −70°C until further use. Cell lysis and protein solubilization was conducted as previously reported. In brief, cells were incubated 15 min at 90°C in 2 % sodium lauroyl sarcosinate (SLS) and 100 mM ammonium bicarbonate and then sonicated for 15 seconds (Vial Tweeter, Hielscher). Soluble proteins were reduced via incubation with 5 mM Tris (2-carboxy-ethyl) phosphine (TCEP) at 90°C for 15 min, followed by alkylation with 10 mM iodoacetamide for 15 minutes at 25°C. Protein concentrations were quantified via a BCA protein assay kit (Thermo Fisher Scientific). 50 μg of protein were digested with 1 μg trypsine (Promega) in 0.25 % SLS (diluted with 100mM ammonium bicarbonate) overnight at 30°C. Following SLS removal via centrifugation, trifluoroacetic acid (TFA) was added to a final concentration of 1.5 % and samples incubated at room temperature for 10 minutes. The supernatant was purified using C18 Micro Spin Columns (Harvard Apparatus) according to the manufacturer’s instructions, dried and resuspended in 0.1 % TFA. Peptide mixtures were then analyzed using liquid chromatography-mass spectrometry carried out on an Exploris 480 instrument connected to an Ultimate 3000 RSLC nano with a Proflow upgrade and a nanospray flex ion source (all Thermo Scientific). Peptide separation was performed on a reverse phase HPLC column (75 μm x 42 cm) packed in-house with C18 resin (2.4 μm, Dr. Maisch). The following separating gradient was used: 94 % solvent A (0.15 % formic acid) and 6 % solvent B (99.85 % acetonitrile, 0.15 % formic acid) to 25 % solvent B over 95 minutes and to 35 % B for additional 25 minutes at a flow rate of 300 nl/min. DIA-MS acquisition method was adapted from Bekker-Jensen et al.^31^. In short, Spray voltage was set to 2.0 kV, funnel RF level at 45, and heated capillary temperature at 275 °C. For DIA experiments full MS resolutions were set to 120.000 at m/z 200 and full MS AGC target was 300 % with an IT of 50 ms. Mass range was set to 350–1400. AGC target value for fragment spectra was set at 3000 %. 49 windows of 15 Da were used with an overlap of 1 Da. Resolution was set to 15,000 and IT to 22 ms. Stepped HCD collision energy of 25, 27.5, 30 % was used. MS1 data was acquired in profile, MS2 DIA data in centroid mode.

Analysis of DIA data was performed using DIA-NN version 1.8^32^, using the UniProt protein database from *Cupriavidus necator H16* and added sequences for formate-THF ligase (*ftfL*, UniProt: Q83WS0), 5,10-methenyl-THF cyclohydrolase (*fchA*, UniProt: Q49135) and 5,10-methylene-THF dehydrogenase (*mtdA*, UniProt: P55818) from *Methylorubrum extorquens AM1*, RK2 plasmid replication protein (trfA, UniProt: P07676), pBBR1 replication protein (pSEVA331 derived AA sequence), aminoglycoside 3’-phosphotransferase (aphA1, Uniprot: P00551) and chloramphenicol acetyltransferase (cat, Uniprot: P62580). Full tryptic digest was allowed with three missed cleavage sites, and oxidized methionines and carbamidomethylated cysteines. Match between runs and remove likely interferences were enabled. The neural network classifier was set to the single-pass mode, and protein inference was based on genes. Quantification strategy was set to any LC (high accuracy). Cross-run normalization was set to RT-dependent. Library generation was set to smart profiling. DIA-NN outputs were further evaluated using a SafeQuant version modified to process DIA-NN outputs^33,34^.

The different metabolic groups were defined as follows. “rGlyP-C1” is comprised of FtfL, Fch and MtdA, “rGlyP-C2” of GcvT1HP and “rGlyP-C3” of SdaA and GlyA. The “plasmid backbone” proteome is composed of TrfA, AphA1, pBBR Rep and Cat. “CBB cycle” proteins contain when applicable CbbL1, CbbL2, CbbS, CbbS2, CfxP, CbxXC, CbbYC, CbbAC, CbbAP, FbP2, Fbp3, Rpe1, Rpe2, CbxXP, CbbTC, CbbTP, CbbZC, CbbZP, CbbKC, CbbKP, CbbGC and CbbGP. The “hydrogen utilization” proteome contains the proteome fractions of HoxA, HoxB, HoxC, HoxF, HoxG, HoxH, HoxI, HoxK, HoxL, HoxM, HoxN, HoxO, HoxQ, HoxR, HoxU, HoxV, HoxW, HoxY, HoxZ, HypA, HypB, HypB2, HypC, HypD, HypE, HypF1, HypF2 and HypX. “Formate utilization” is comprised of the proteins FdsD, FdsA, FdsB, FdsG, FdsR, FdoI, FdoH, FdoG, FdhA1, FdhA2, FdhB1, FdhC, FdhD, FdhD1, FdhD2, FdhE, FdwA, FdwB and CbbB. All other detected proteins (~3500) make up the “other” category.

## Supplementary Texts

### Supplementary Text S1: Integration and sequencing of various C1-operon integration strains

The strain CRG4.5 (CRG4 cured of pC1) was, as expected, unable to grow on minimal media supplemented with formate as sole carbon and energy source (Fig. S3a). Next, we used this strain to randomly integrate the C1 module using transposon integration vectors with promoter variants of different strengths controlling the synthetic C1-operon^18^,^35^. Following conjugation of the transposon vectors for each of the different promoters, the cell populations were transferred to formate minimal media to select for formatotrophic growth (Fig. 1b). Populations were repeatedly inoculated into fresh medium until the growth rate of the cell population ceased to increase (Fig. S4a). After 10 passages, isolated fast-growing clones with different promoter combinations showed similar growth rates and were characterized by sequencing (Fig. S5a, Table S1).

In these analyses, we found one of these clones (from here on CRG5) to contain two separate insertions of the same P_14_-C1 operon, indicating it may have selected for optimal expression levels by integrating two copies of the C1 operon (Fig. S6a, Table S1). The first insertion interrupted the gene encoding a methyl-cis-aconitate hydratase (*acnM*) involved in the 2-methylcitrate cycle for growth on propionate^36^, which is redundant for formatotrophic growth conditions. The second insertion was found upstream of the gene encoding a phasin protein in *C. necator (phaP1*), which is involved in encapsulating the storage polymer polyhydroxybutyrate (PHB). *phaP1* is highly expressed in PHB producing conditions and can make up 5 % of the total protein in *C. necator*^37^. However, the *C. necator* production platform strains used in this study are incapable of PHB formation due to a *phaC1* knockout, making the *phaP1* expression redundant and possibly making it a good locus for heterologous expression as the C1 operon may benefit from additional expression by read-through from the native *phaP1* promoter.

Sequencing of the CRG5 strains with other promoter variants than P_14_ showed more insertions in the *phaP1* locus. Clones with the C1 module under expression of the promoters P_3_, P_4_ and P_14_ (2^nd^ clone) all had the operon inserted in the *phaP1* locus 98 nt upstream of the start codon, further highlighting the relevance of this locus (Table. S3). Overall, the sequencing results from different clones indicate that the weaker promoters (such as P_14_) were integrated in this high expression locus to support fast formatotrophic growth (Table S3). The C1 operon under control of the strongest tested promoter (P_2_) however inserted the transposon in the *H16_A3661 (H16_B0502*) locus but also deleted 189 nt of the *phaP1* promoter region (+123 nt) (Table S3). Also, this was the only sequenced clone where the C1 operon inserted anti-sense with respect to the surrounding ORF or promoter directionality. This indicated that expression from the strong P_2_ promoter did not require additional expression via read-through from the *phaP1* promoter. The P_PhaC_-C1 transposon integrated at position 590 nt in the short chain dehydrogenase gene *H16_A0258*, possibly associated with PHB binding^38^, and a mutation locus also later found in CRG6. The strain with the weakest constitutive promoter (P_14_) and two C1 operon insertions was further characterized in detail (referred to as CRG5 in the main text).

### Supplementary Text S2: Integration of the C3 operon and the DadA6 route

To investigate if the growth rate and yield of the rGlyP strains could be further optimized, we next aimed at integrating module 3 using the workflow described above (Fig. 1b). We cured the CRG5 strain of the last remaining plasmid (pC3). Notably, the created CRG5.5 strain was still able to grow on formate, albeit at a much slower growth rate and biomass yield (Fig. S3a,b). This is in line with our previous study, where we identified glycine oxidase DadA6 as the native glycine assimilation route, which could be used as an alternative to the plasmid-encoded C3 module (Fig. S1a). The native route wastefully oxidizes glycine to glyoxylate, which is further channeled into phospho-glycerate and central metabolism via the glycerate shunt. The glycine oxidation route is energetically on par with the CBB Cycle, limiting the biomass yield of the CRG5.5 strain^18,39^.

To re-establish the superior serine route, we transformed CRG5.5 with another set of transposon integration vectors that contained native *C. necator sdaA* and *glyA* genes downstream of another small library of constitutive promoters and selected for formatotrophic growth (Fig. S4b). After 10 total passages on formate minimal media, several similarly fast-growing strains (CRG6) were isolated. Among the different constructs, the CRG6 lineage containing the P_3_-C3 module seemed to perform best (Fig. S5b). Sequencing revealed that in one of the strains the P_3_-C3 operon interrupted the *H16_B1976* gene, encoding a putative *tonB*-dependent outer membrane receptor (Table S2). In the other sequenced strain the P_3_-C3 operon was inserted into the *prpB* gene located on chromosome 1. PrpB, together with AcnM is part of propionate metabolism in *C. necator* (Table S3)^36^. The CRG6 clone with the P_3_-C3 operon integrated in the *H16_B1976* gene grew best and was further investigated (referred to as CRG6 further).

To suppress the glycine oxidation route in CRG6 that might compete with the rGlyP, we deleted *dadA6*. However, the doubling time and yield of the CRG6 Δ*dadA6* strain were not significantly altered (Fig. S3a,b), indicating that the glycine oxidation pathway did not play a major role for growth of CRG6 on formate.

### Supplementary Text S3: In-depth proteomic analysis of formatotrophic strains

First, we compared the expression of the rGlyP genes in the CRG4 and CRG6 strains (Fig. 3a, S7a). Here we found the genomic expression of module 1 from strain CRG6 to have almost reached the expression of the same operon from plasmid in CRG4. As both the plasmid-based and genomic C1 operons are controlled by the same (weak) P_14_ promoter we attribute the consistent high expression from the genome to the duplicate integration in the genome and read-through expression of strong *phaP1* promoter. Expression of module 2 (GCV) remained basically unchanged from CRG4 to CRG6, which was expected, as this module was already genome-integrated previously^18^. The genomic insertion of module 3 of the CRG6 strain however decreased expression compared to the CRG4 strain by over one order of magnitude, possibly indicating that this operon was expressed too strongly from plasmid in the CRG4 strain (Fig. 3a, S7a).

Next, we investigated the effect of the transposon insertions and mutations on their insertion loci. Expression of genes in the C1-operon integration loci were decreased (*phaP1* decreased 150-fold and *acnM* by a factor of 14) (Fig. S7a). The short chain dehydrogenase H16_A0258, which contains a 1 bp deletion in strain CRG6 (Table S2) and was also found to be a target of the P_PhaC_-C1 transposition event was also downregulated by a factor of 28. The protein encoded by the gene *H16_B1976* and target of the gP_3_-C3 transposon insertion was not up or down regulated significantly, possible reflecting the neutrality of this locus in terms of expression and deletion for the cell.

We then compared the proteomes of CRG6 to CRG4 beyond rGlyP and mutation loci (Fig. S7a). Interestingly, among the most upregulated proteins in CRG6 compared to CRG4 is 2-amino-3-ketobutyrate CoA ligase (Kbl), which is upregulated by a factor of 26. Kbl, together with threonine dehydrogenase (Tdh) catalyzes the formation of threonine from glycine and acetyl-CoA. Tdh however was not significantly up or down regulated in CRG6, and acetyl-CoA could only be wastefully generated from pyruvate via rGlyP and pyruvate dehydrogenase. This leaves an unclear picture of the relevance of the Kbl upregulation in CRG6. Glutamate dehydrogenase 1 (Gdh1) seems to be slightly upregulated (~4 fold) in CRG6 over CRG4. This however, is due to a 13-fold downregulation of Gdh1 in CRG4 compared to the CBB cycle strain. PHB de-polymerase (PhaZ6), as well as formate dehydrogenase subunits FdhA1 and FdhC are downregulated by ~2-fold, which possibly further relates to the biomass yield gain of CRG6 over CRG4. Overall, proteins associated with drug efflux, *tonB*-dependent receptors, ferric uptake and L-leucine/L-phenylalanine ABC transporters where upregulated in CRG6 relative to CRG4 (Fig. S7a).

Next, we used measured protein intensities to allow approximate quantification of the total proteome fractions allocated to different metabolic tasks (Fig. 3b). We found the rGlyP to make up 25 % of the quantified proteome in CRG4 and around 14 % in CRG6. In strains CRG4 and CRG6 the C1 module accounted for 10 % and 8 % respectively. Expression of module 2 in CRG4 in CRG6 made up 5 %, while module 3 contributed to a fraction of 9 % in CRG4 and only 0.8 % in CRG6. The proteome fraction associated with Rubisco in the CBB strain made up ~1.9 %, with all of the CBB cycle enzymes together only accounting for 6 % of the proteome. Despite Rubiscos famously high protein content in higher plants our findings are in line with previous studies of bacterial Rubisco protein abundance^40^. The Rubisco protein concentration is further negatively correlated with elevated CO_2_ conditions and all strains were grown at high (10 %) CO_2_^41^. We find the hydrogen metabolism proteins to make up a fraction of 3 % in the CBB strain. However, the CBB cycle and hydrogen readiness constitute only 0.01 % and 0.1 % respectively of the total quantified proteome in the rGlyP strains (Fig. 3b).

Last, we compared the proteomes of CRG4 and CRG6 to the CBB strain (Fig. S7b,c). Here we found several global proteome changes that we could partially relate to mutations already present in the CRG4 lineage. In all rGlyP dependent strains the CBB cycle is downregulated by two orders of magnitude, which was expected due to the deletion of the transcriptional activator *cbbR*^18^. Also, the hydrogen utilization proteome is downregulated by a similar degree in line with the *hoxA* mutation already present in the CRG4 strain lineage. This indicated that the CRG4 and CRG6 strains lost their “readiness”^23^ to use hydrogen as substrate, which is indeed redundant for formatotrophic growth. We further found propionyl-CoA transferase Pct to be upregulated by factor of 16, possibly related to mutations found in the 2-methylcitrate cycle genes *acnM* and *prpB*. Since the rGlyP is connected to central metabolism at the level of pyruvate, we expected more gluconeogenic flux compared to the CBB strains. While this was not reflected by significantly altered expression of the genes encoding PEP synthetase, Pyruvate carboxylase and PEP carboxykinase, we did observe pyruvate kinase 3 to be downregulated by a factor of 16 in both CRG4 and CRG6 (Fig. S7b,c).

### Supplementary Text S4: FDH inhibition experiment

We confirmed that sFDH levels in CRG6 were optimized by titration experiments in which we supplied either the sFDH inhibitor tungsten or its required metal co-factor molybdenum (Fig. S8). While these experiments did not show further increases in the biomass yield of CRG6 at any FDH inhibition or ‘induction’ level, slight sFDH inhibition (10nM tungstate) in CRG4 increased the yield to the highest level observed so far for this strain. This indicated that in CRG4 formate flux had not been fully optimized, while in CRG6 formate oxidation and assimilation were apparently balanced.

## Supplementary Figures

**Fig. S1:**
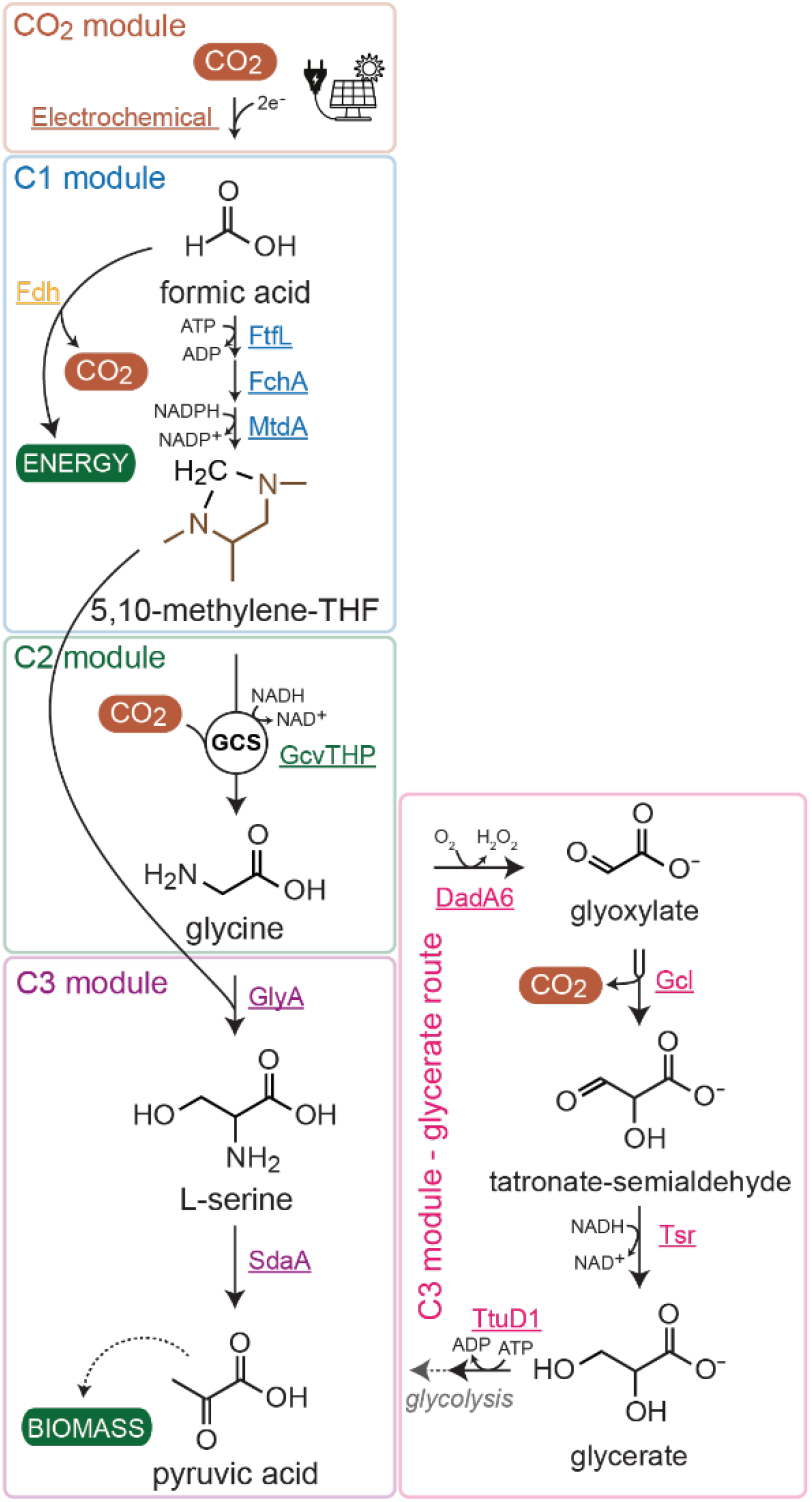
The architecture of the reductive glycine pathway (rGlyP). The rGlyP serine variant is compared to the glycine oxidation route via *dadA6* natively present in *C. necator*. This representation of the rGlyP also shows the proposed formate production from CO_2_ using renewable energy and electrochemical reduction (CO_2_ module). Formate is then activated and reduced to 5,10-methylene-THF via the 3 heterologous enzymes from *M. extorquens* formate-THF ligase (FtfL), 5,10-methenyl-THFcyclohydrolase (FchA) and 5,10-methylene-THF dehydrogenase (MtdA) (C1 module). Next, methylene-THF is aminated, condensed with CO_2_ and reduced to glycine by the glycine cleavage system (gcvT1HP) operating in the reductive direction (C2 module). Last, glycine is converted to pyruvate via one of two depicted routes (C3 module). Glycine is either condensed with another methylene-THF via serine hydroxymethyltransferase (GlyA) to yield L-serine, which is then dehydrated by serine deaminase (SdaA) to yield pyruvate. Alternatively, glycine can be oxidized to glyoxylate by glycine oxidase (DadA6), glyoxylate then is metabolized via the glycerate shunt to glycerate-phosphate via glyoxylate carboligase (Gcl), tatronate-semialdehyde reductase (Tsr) and glycerate kinase (TtuD1). Pyruvate then is generated via glycolysis.

**Fig. S2:**
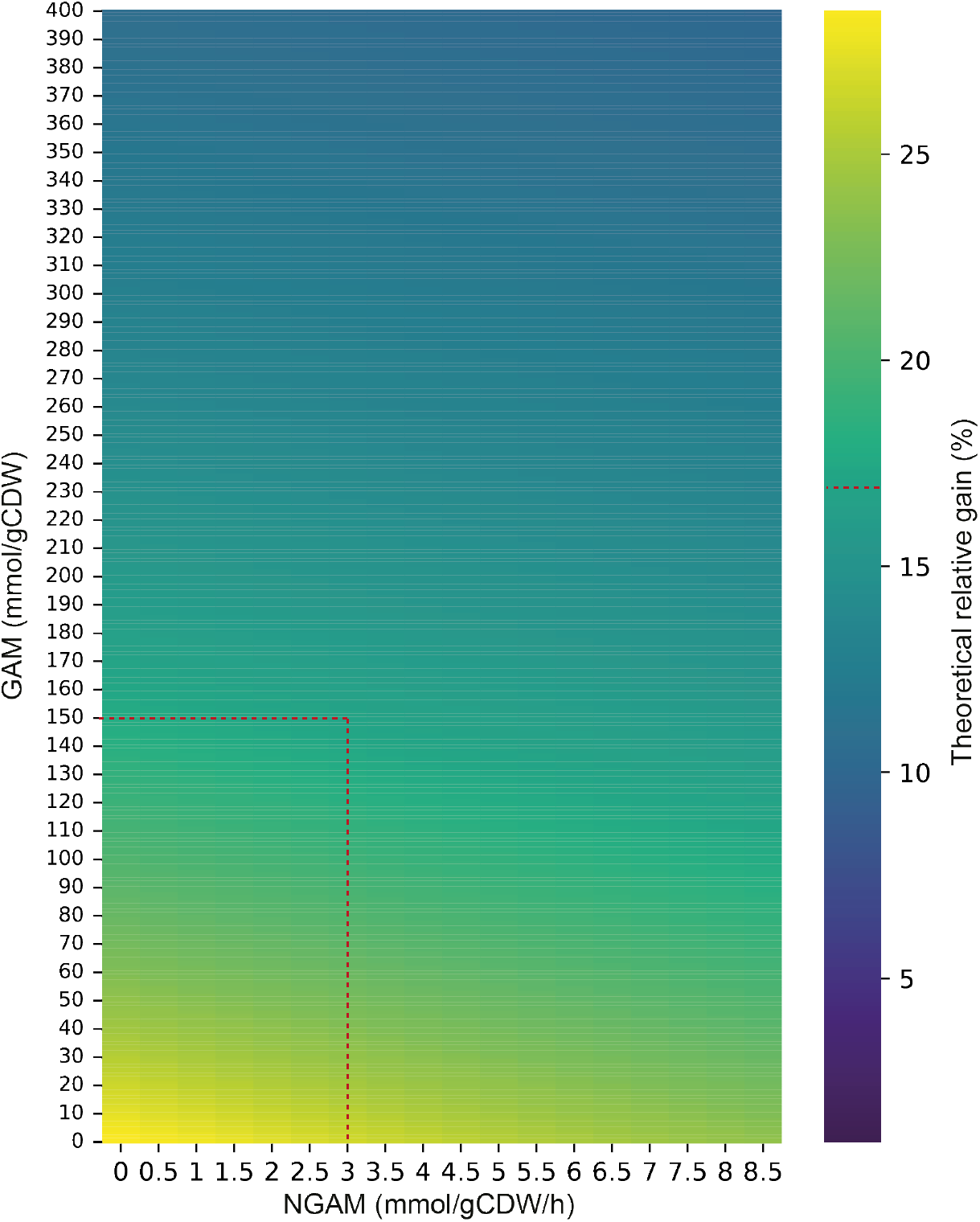
Predicted biomass yield increase of the rGlyP over CBB. Theoretical relative gain (%) in biomass yield of the reductive glycine pathway (rGlyP) over the CBB cycle in *Cupriavidus necator* at a doubling time of 4 h. Red dashed lines indicate the theoretical relative gain of *~*17 % at GAM=150 mmol/gDW and NGAM=3 mmol/gDW/h (parameters derived from Jahn et al^31^).

**Fig. S3:**
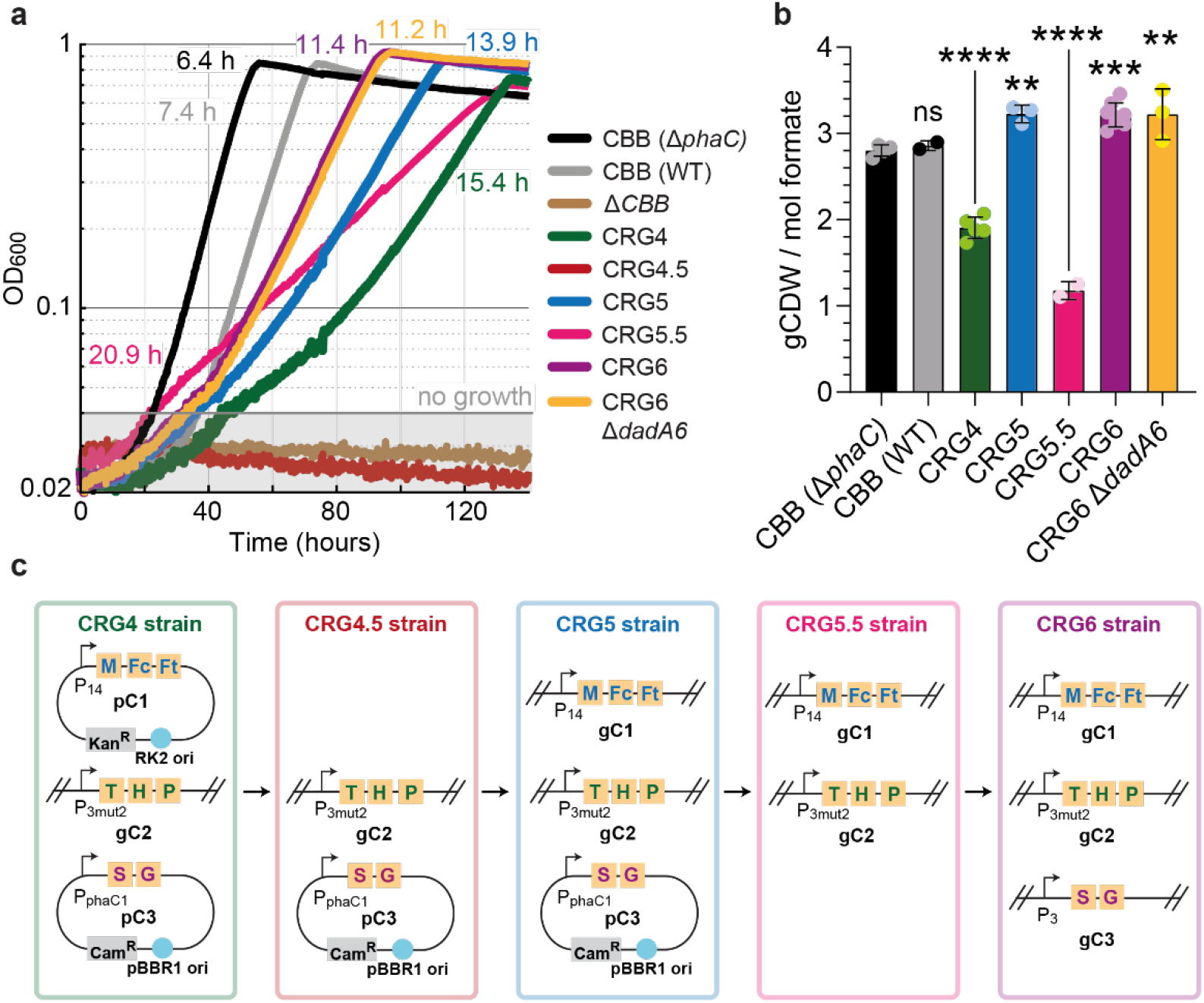
Overview of strains and their phenotypes. **a**, Growth of various *C. necator* strains used in this study in M9 minimal media supplemented with 80 mM formate and 100 mM bicarbonate with 10 % CO_2_ in the headspace. The doubling time in hours (h) of the strains is presented in the designated strain color. Curves depict the mean of 2-3 technical replicates and are representative of 3 experiments conducted in the same conditions to ensure reproducibility. Less than one doubling of the bacterial strains was considered no growth and indicated as such via an overlayed shaded box. **b**, Biomass yields in gram cell dry weight per mol formate consumed of *C. necator* H16 CBB (Δ*phaC*), CBB (WT), CRG4, CRG5, CRG5.5, CRG6 and CRG6 Δ*dadA6* flask cultures (n ≥ 2). Yield data were obtained from 4 and 8 biological replicates of 2 and 3 experiments respectively for Δ*phaC* and CRG6. WT values correspond to one experiment with two biological replicates. CRG4 values correspond to 5 biological replicates from 2 experiments. CRG5, CRG5.5 and CRG6 Δ*dadA6* consist of 3, 2 and 3 biological replicates respectively from one experiment. All data points and their mean are shown. Error bars indicate standard deviation. Significant differences of the individual strains relative to the CBB (Δ*phaC*) control strain are indicated. Two asterisks (**) represent a p-value < 0.01, three asterisks (***) a p-value < 0.001 and 4 asterisks (****) a p-value < 0.0001. **c**, Simplified strain overview and localization of the reductive glycine pathway modules (C1-C3) in the different strains used in this study. Plasmid-based expression (p) and genomic expression (g) are indicated.

**Fig. S4:**
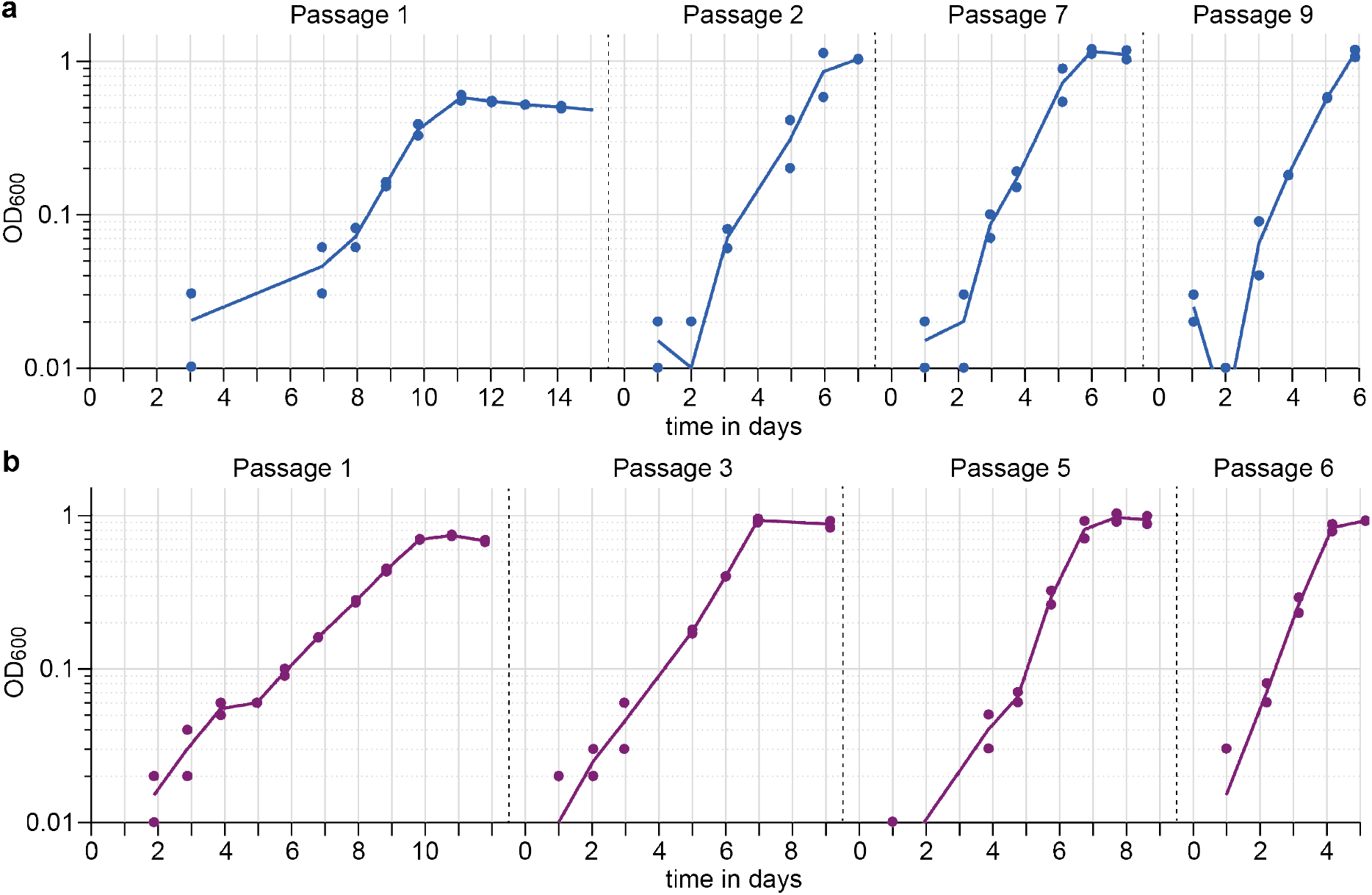
Creation of CRG5 and CRG6 strains. Liquid selection test tube experiment in M9, 80 mM formate, 100 mM HCO_3_ minimal media for creation of the CRG5 gP_14_-C1 (**a**) and CRG6 gP_3_-C3 strains (**b**). Following transposition of module 1 or module 3 of the rGlyP and upon reaching early stationary phase cells were re-inoculated into fresh formate media. Dots represent the measured glass tube OD_600_ values of two biological replicates per promoter operon transposon insertion. The lines correspond to their averages. 4 passages are depicted and are indicative of the improved growth over the 10 passages.

**Fig. S5:**
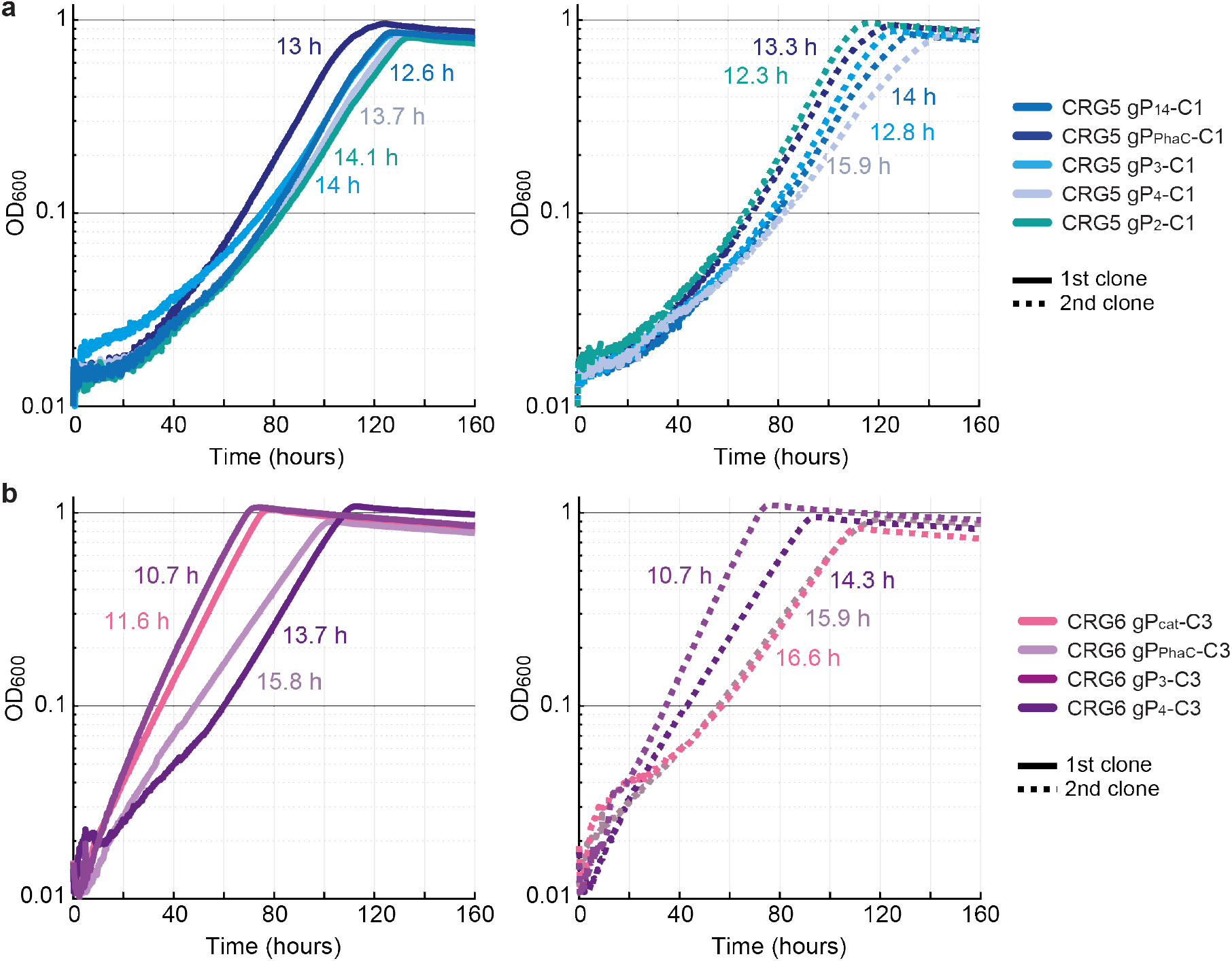
Characterisation of CRG5 and CRG6 lineages with different promoters. Growth profiles of isolated single clones after the liquid selection experiment of (**a**) CRG5 and (**b**) CRG6 strains with different promoter C1 and C3 module transposon insertions in M9, 80 mM formate, 100 mM HCO_3_ minimal media. Promoters P_14_, P_PhaC_, P_3_, P_4_ and P_2_ were used to control the C1 operon, while the promoters Pcat, P_PhaC_, P_3_ and P_4_ were used to control the C3 operon. Two individually transposed lineages are shown per promotor. A solid line indicates the first clone and a dotted line the second. Doubling time in hours is given in the respective strain color beside the growth curve. Curves are means of 3 or 4 technical replicates for CRG5 and CRG6 respectively and representative of 3 experiments conducted in the same conditions. 1^st^ clone strains CRG5 gP_14_-C1 and CRG6 gP_3_-C3 were the focus of this study and have the corresponding color shade associated with them in the main text figures.

**Fig. S6:**
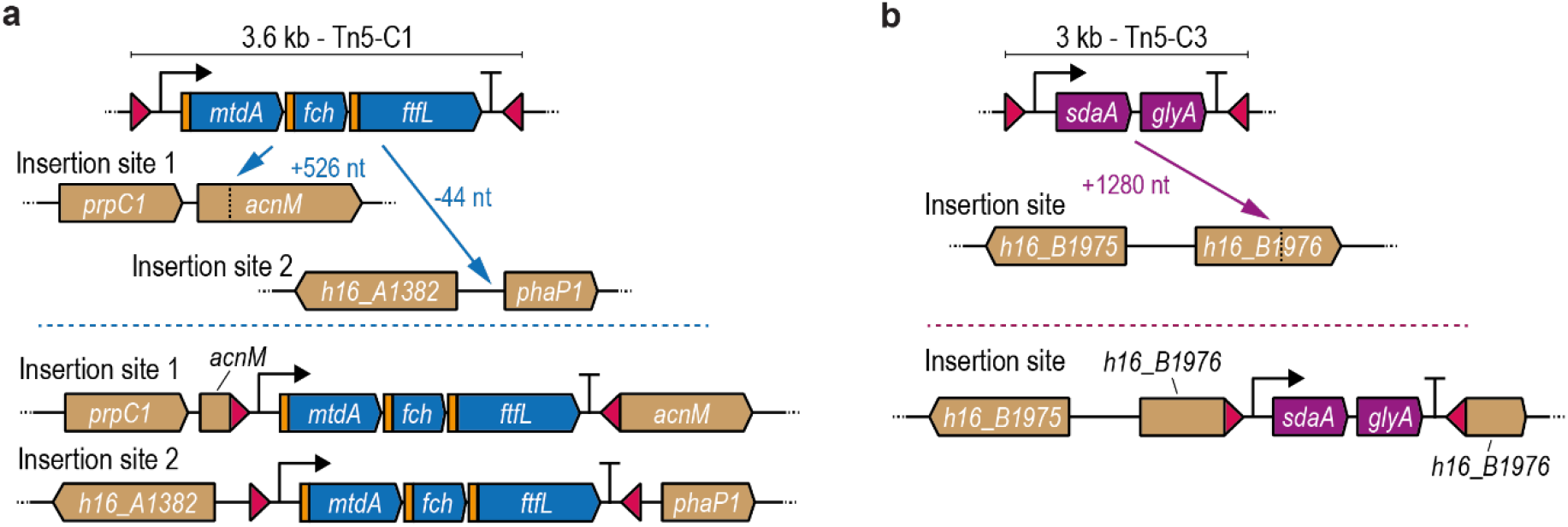
Transposon insertion loci of the genomic rGlyP modules in this study. **a**, the CRG5 lineage carries two insertions of the Tn5-P_14_-C1 construct. One in the *acnM* gene and one upstream of the *phaP1* gene. **b**, the transposon expressing the P_3_-C3 module inserted in the gene *H16_B1976* in strain CRG6.

**Fig. S7:**
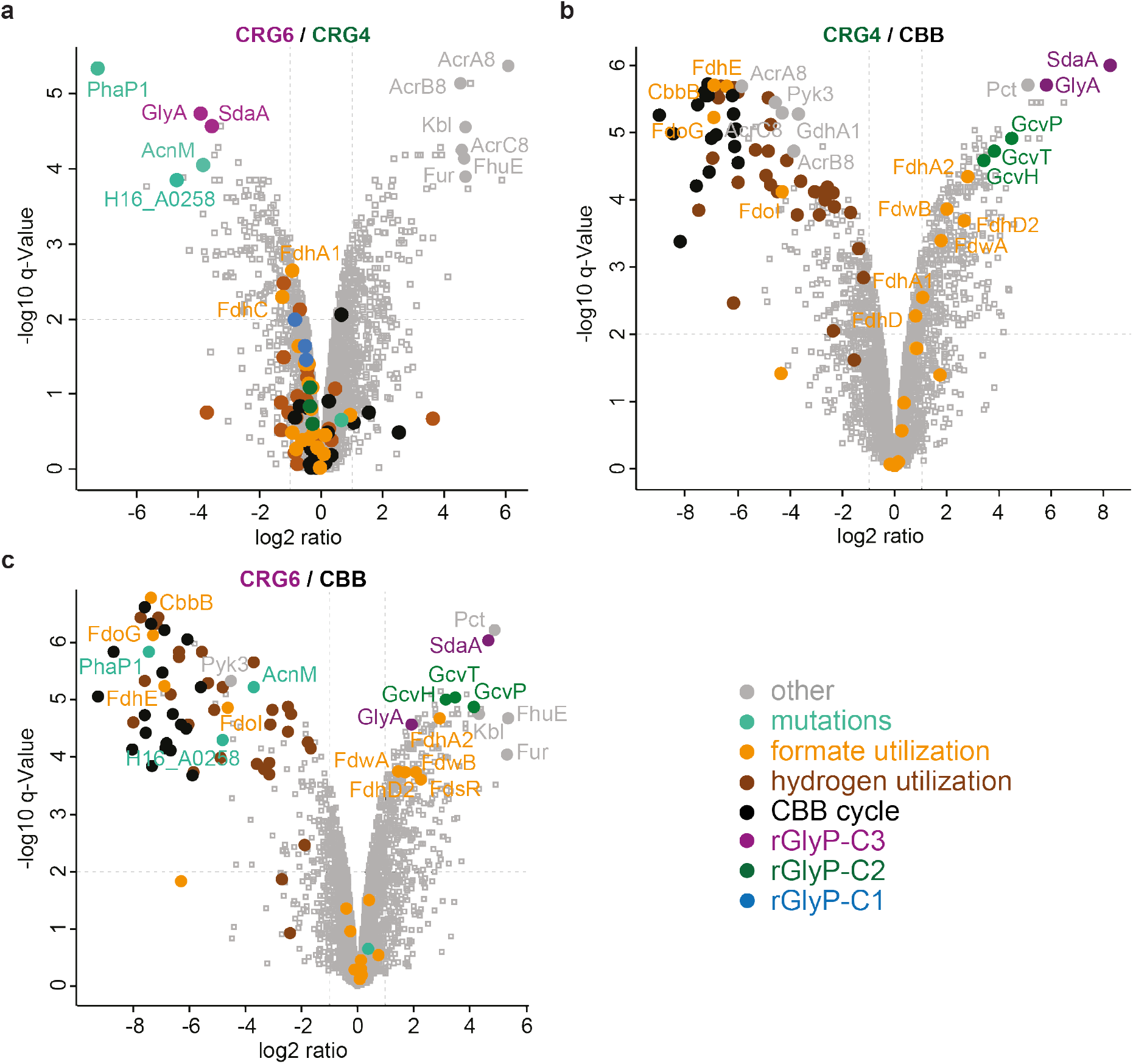
Proteomic comparison of rGlyP and CBB strains. Global changes in the proteomes of (**a**) CRG4 and (**b**) CRG6 compared to the CBB strain and of CRG6 to CRG4 (**c**). All strains were grown in formate minimal media in shake flasks. Log2-fold changes in protein intensity are plotted against significance (-log 10 q-Values). Relevant proteins are labeled, depicted as a filled-out circles and colored according to their metabolic task. rGlyP proteins associated with module 1 are shown in blue, module 2 in green and module 3 in purple. CBB cycle and hydrogen utilization proteome are depicted in black and brown respectively. Different formate utilization proteins are highlighted in orange. Loci hit by transposon insertions and frequently observed with mutations such as *H16_A0258* are shown in turquois. The most up and downregulated proteins if not already named, are colored in grey. All other proteins are not labeled, depicted as empty squares and colored grey.

**Fig. S8:**
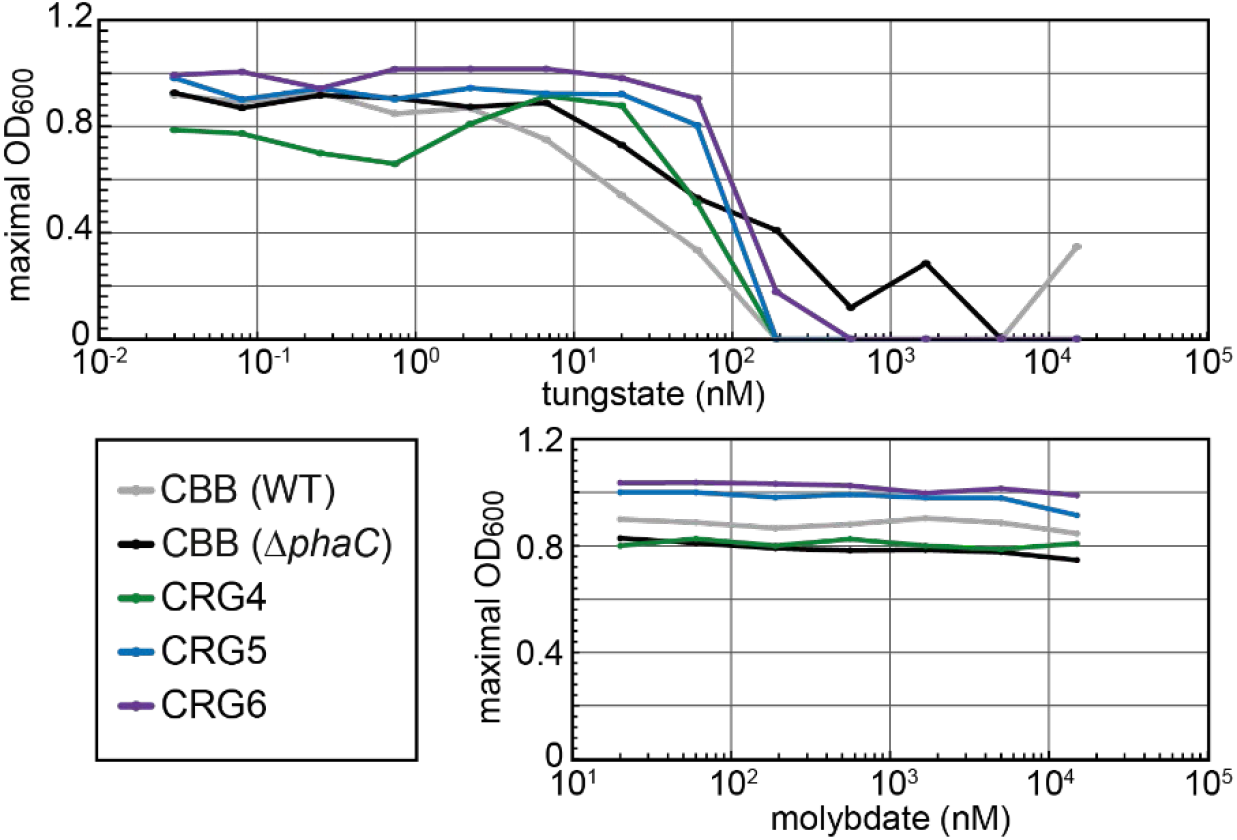
Effect of sFDH inhibition on biomass yield (OD_600_). *C. necator H16* CBB (WT), CBB (Δ*phaC*), CRG4, CRG5 and CRG6 strains were grown in M9, 80mM formate, 100mM HCO_3_ minimal media in 96-well plates. Cells were either supplemented with varying concentrations of sodium tungstate to inhibit sFDH or sodium molybdate to potentially further increase sFDH activity with the appropriate metal ion.

## Supplementary tables

**Table S1:** Mutations of CRG5 gP_14_-C1 compared to CRG4.5. Chromosome is abbreviated as Chr.

| Chr. | position | mutation | annotation | gene | description |
| --- | --- | --- | --- | --- | --- |
| 1 | 1,499,624 | Tn5-P<sub>14</sub>-C1 insertion | Insertion 44 nt upstream of start ATG | *phaP1* | phasin (PHA-granule associated protein) |
| 1 | 2,072,499 | Tn5-P<sub>14</sub>-C1 insertion | Insertion after I175 (525/2613 nt) | *acnM* | methyl-cis-aconitic acid hydratase/Aconitate hydratase |
| 2 | 222,178 | (AGGCCG CGCGGC) 1→2 | coding (1489/3063 nt) | *sbcC* | DNA repair exonuclease, SbcC |
| 2 | 926,039 | G→C | L32V (CTC→GTC) | *B0818* | transcriptional regulator, XRE-family |

**Table S2:** Mutations of CRG6 gP_3_-C3 compared to CRG5.5. Chromosome is abbreviated as Chr.

| Chr. | position | mutation | annotation | gene | description |
| --- | --- | --- | --- | --- | --- |
| 1 | 268,712 | Δ1 bp | coding (10/855 nt) | *A0258* | short chain dehydrogenase |
| 1 | 3,461,712 | +G | coding (823/969 nt) | *A3204* | transcriptional regulator, LysR-family |
| 2 | 159,389 | +G | Insertion 22 nt upstream of start ATG | *B0138* | transcriptional regulator, IclR-family/conserved hypothetical transmembrane protein |
| 2 | 2,234,079 | Tn5-P<sub>3</sub>-C3 insertion | Insertion at (1279/1935) | *B1976* | outer membrane receptor, TonB dependent |

**Table S3:** Tn5 promoter C1 and C3 operon insertion sites of CRG5 and CRG6 clones. *location on chromosome (Chr.) 1 or 2 of the P_2_-C1 insertion is unclear, as it inserted in a large duplicated region with identical sequence on chromosome 1 and 2.

| Strain | Chr. | Position (nt) | Insertion locus | Direction |
| --- | --- | --- | --- | --- |
| CRG5 gP<sub>14</sub>-C1 | 1 | 1,499,624 | *phaP1/H16_A1382* | sense |
|  | 1 | 2,072,499 | *acnM* | sense |
| CRG5 gP<sub>PhaC</sub>-C1 | 1 | 269,292 | *h16_A0258* | sense |
| CRG5 gP<sub>3</sub>-C1 | 1 | 1,499,668 | *phaP1/H16_A1382* | sense |
| CRG5 gP<sub>4</sub>-C1 | 1 | 1,499,668 | *phaP1/H16_A1382* | sense |
| CRG5 gP<sub>2</sub>-C1 | 1 | 3,942,813 | *H16_A3661 \*or H16_B0502* | anti-sense |
| CRG5 gP<sub>14</sub>-C1-2 | 1 | 1,499,668 | *phaP1/H16_A1382* | sense |
| CRG6 gP<sub>3</sub>-C3 | 2 | 2,234,079 | *H16_B1976* | sense |
| CRG6 gP<sub>3</sub>-C3-2 | 1 | 2,070,024 | *prpB* | sense |

